# StereoSiTE: A framework to spatially and quantitatively profile the cellular neighborhood organized iTME

**DOI:** 10.1101/2022.12.31.522366

**Authors:** Xing Liu, Chi Qu, Chuandong Liu, Na Zhu, Huaqiang Huang, Fei Teng, Caili Huang, Bingying Luo, Xuanzhu Liu, Min Xie, Feng Xi, Mei Li, Liang Wu, Yuxiang Li, Ao Chen, Xun Xu, Sha Liao, Jiajun Zhang

## Abstract

With emerging of Spatial Transcriptomics (ST) technology, a powerful algorithmic framework to quantitatively evaluate the active cell-cell interactions in the bio-function associated iTME unit will pave the ways to understand the mechanism underlying tumor biology. This study provides the StereoSiTE incorporating open source bioinformatics tools with the self-developed algorithm, SCII, to dissect a cellular neighborhood (CN) organized iTME based on cellular compositions, and to accurately infer the functional cell-cell communications with quantitatively defined interaction intensity in ST data. We applied StereoSiTE to deeply decode ST data of the xenograft models receiving immunoagonist. Results demonstrated that the neutrophils dominated CN5 might attribute to iTME remodeling after treatment. To be noted, SCII analyzed the spatially resolved interaction intensity inferring a neutrophil leading communication network which was proved to actively function by analysis of Transcriptional Factor Regulon and Protein-Protein Interaction. Altogether, StereoSiTE is a promising framework for ST data to spatially reveal tumoribiology mechanisms.

## Introduction

Cell-cell communication by molecular interaction within immune tumor microenvironment (iTME) is closely related to tumorigenesis, progression, and treatment response, which provides important targets for diagnosis, prognosis, disease monitoring, drug design, etc^1^. The advanced spatial transcriptomic (ST) technology provides opportunities to better elucidate physiological and pathological progress by observing the comprehensive nature of transcriptome in spatial^2, 3^. The expression profile with spatial coordination empowers the ability to reveal the landscape of iTME and to dissect the cell-cell communication in pathogenesis associated iTME region^3^. However, how to precisely find pathogenesis associated iTME, quantitatively infer cell-cell communications and reasonably utilize spatial information, remain to be the significant challenges. Most open source software for cell-cell communication analysis are designed based on either gene expression^4, 5^ or biological distance, without considering the interactable cell-cell distance^6–8^. Comprehensively understanding the importance of spatial information needs a well-designed framework composed of powerful algorithms to decode spatial data.

Here, we present StereoSiTE, an analytical framework for comprehensive depiction of landscape of iTME which is defined by Cellular Neighborhood (CN)^9^ and dissection of spatial cell interaction intensity (SCII). CNs are defined by the cellular composition obtained from deconvolution result and SCII next detects cell-cell communication using both cell space nearest neighbor graph and targeted L-R expression. Moreover, analysis of SCII in CN region of interest can elucidate multicellular communities in correspondent iTME unit, which provides new dimension in evaluating iTME. To exhibit the application scenario, we applied StereoSiTE to analyze ST data of xenograft models receiving immunoagonist treatment and revealed the iTME landscape and spatial cell interactions in functional iTME units, which provided molecular interaction and sequential activities of tumor in response to treatment.

## Result

### StereoSiTE: A novel framework to spatially and quantitatively profile cell-cell communications in cellular neighborhood organized iTME

In this framework (Fig 1A), we first performed cellular neighborhood analysis to investigate the cellular composition based iTME organization. Then, the self-developed SCII was applied to analyze the spatially resolved cell-cell communication to indicate key molecular activities by constructing biological network.

**Figure 1.**
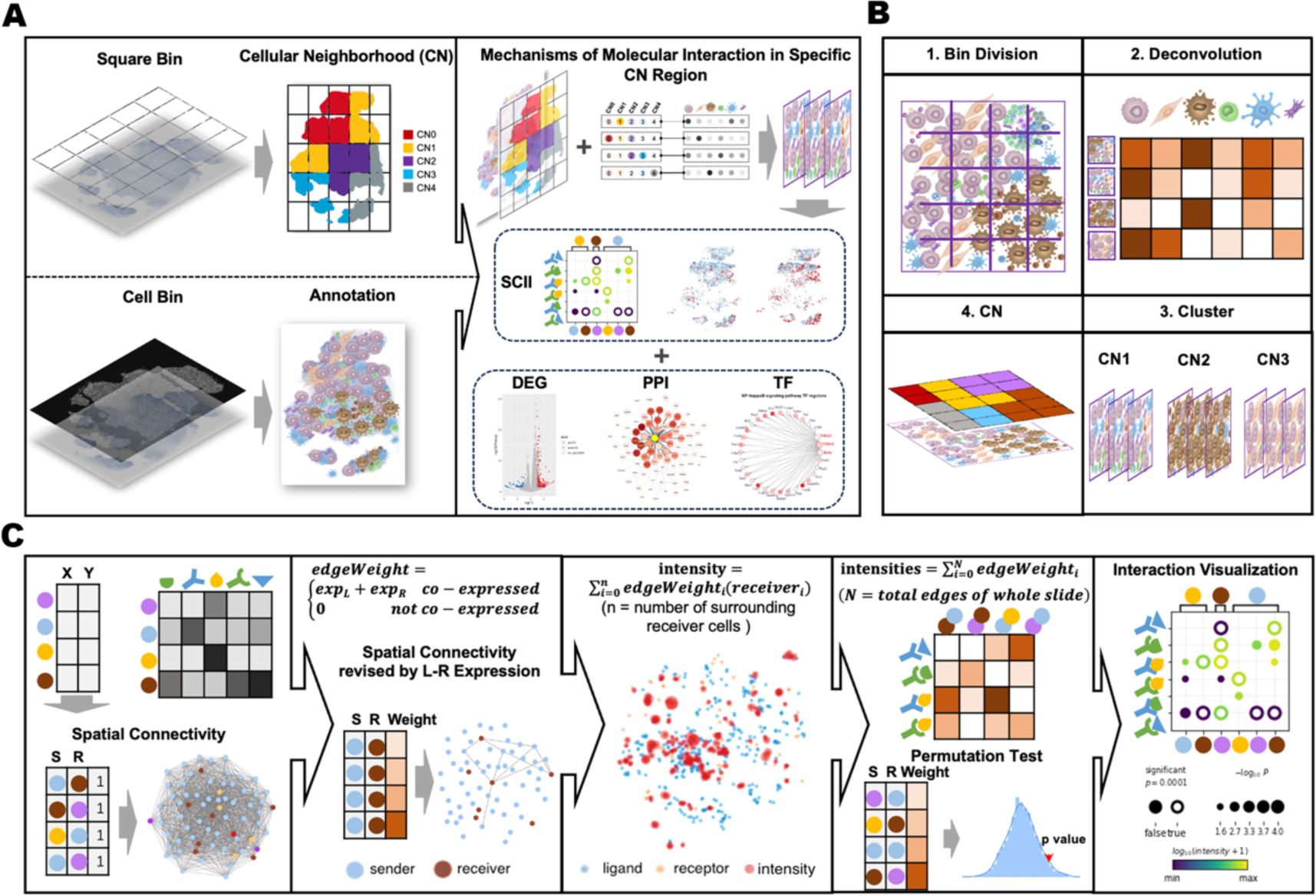
Schematic diagram of StereoSiTE workflow. **A.** Overview of StereoSiTE, including analysis of CN, annotation, tensor deconvolution, SCII and compatible pathway analysis. **B.** Conceptual diagram of CN: 1. Converting the spatial gene expression matrix into binned data matrix. 2. Resolving the cellular compositions by deconvolution (cell2location). 3. Clustering bins into different cellular neighborhoods based on their cellular composition. 4. Mapping the CNs in situ. **C.** Principle of SCII: Firstly, constructed the space nearest neighbor graph based on coordinates of cells; Secondly, revised the space nearest neighbor graph by assigning the connected edges; Thirdly, calculated the interaction intensity; Then, permutation test was used to build a null distribution by shuffling the cell type labels and computed the p value. Finally, the spatial cell interaction intensities and p values were visualized by bubble plots indicating interaction intensity (by color) and p value (by size).

Cellular Neighborhood (CN) was performed on squared binned data based on cellular composition which was resolved by deconvolution method^10^ (cell2location) with single-cell sequencing data as reference. Squared bins with similar cellular composition were clustered together by Leiden^11^. Each cluster can be defined as an iTME unit. To select CNs of interest for functional analysis, we integrated the matrix co-currently covering cellular neighborhoods (CNs) and cell types (CTs) and introduced Tensor^12^ to decompose the module matrix.

To analyze the spatially informed cell to cell interaction, we developed Spatial Cell Interaction Intensity (SCII), which detected actively functional L-R pairs by quantitatively defining interactive intensity with spatial proximity of cells and expression of genes. To improve accuracy, we recommended using spatially resolved data at single cell resolution. Cell segmentation was performed according to published protocol^13^. Then we conducted cell type annotation by cell2location referring to protocol mentioned above. With the annotated single cell resolved data, we constructed the space nearest neighbor graph by connecting cells whose distance within a threshold and revised it by assigning the weight to connected edge with expression. Then, we calculated the local cell interaction intensity in situ by summing up weight of edges linked to each sender cell and the entire interaction intensity of whole slide by summing up weight of all edges. Finally, to count the confidence, we used permutation test to get a null distribution by shifting cell type annotation labels, and p value can be calculated from this null distribution (Fig 1C).

To prove that testable hypotheses can be derived from inferred cell-cell communications, upstream and downstream signaling activities need to be comprehensively analyzed. StereoSiTE also included analysis of Differentially Expressed Genes (DEG), Protein-Protein Interaction (PPI) network and Transcriptional regulatory Factory (TF) network to provide an end-to-end solution for selected L-R pairs induced molecular mechanism of specific iTME unit.

### Decoding iTME into cellular neighborhood (CN) organized units

To verify that CN is a reasonable unit to study iTME, we applied this framework to analyze representative Stereo-seq data from the cancer tissue of xenograft model and analyzed CN based iTME unit. We generated a binned data matrix with bin size of 100 (equal to 50µm*50µm), and decoded the cell-type composition of each bin. The ST data was clustered into 7 different CN clusters (Fig 2A), of each with a unique cell-type composition (Fig 2B). To decode the microenvironment of each CN, we aligned annotated data at single-cell resolution on clustered CN under guidance of coordinate. Fig 2C showed the annotation result in situ, and the proportion of different cell types in this sample was calculated (Fig 2D). Overall, non-immune cells accounted for 77.05%, lymphoid cells accounted for approximately 10.28% and myeloid cells accounted for approximately 12.67%.

**Figure 2.**
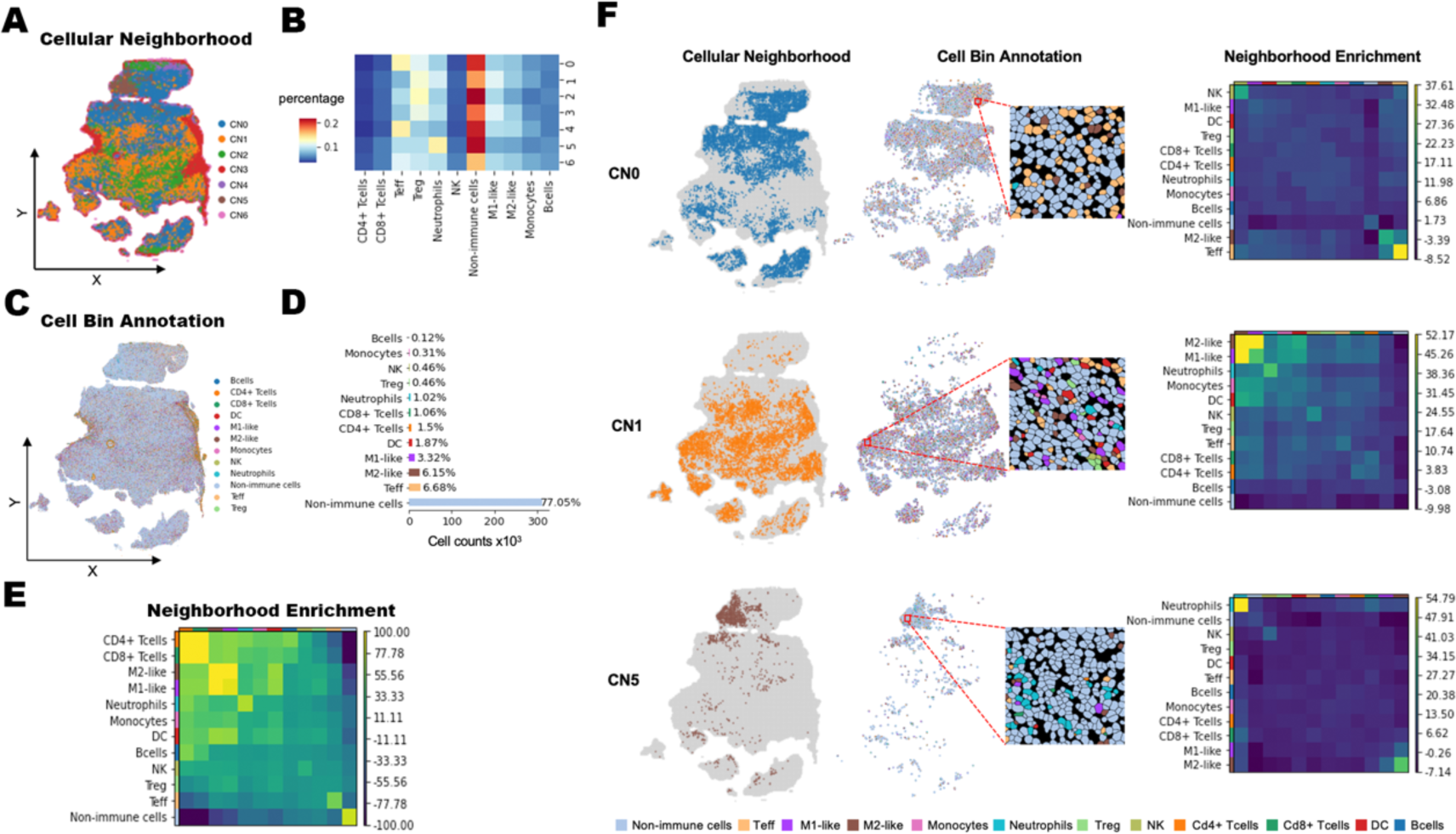
Performance of Cellular Neighborhood (CN) in decoding iTME. **A.** Cellular Neighborhood clustered based on cellular composition. **B.** Heatmap of cellular compositions in individual CN. **C.** Annotation at single cell resolution in situ. Each spot represented a single cell. **D.** The counts and percentages of each cell type, the X-axis represented the cell number. **E.** The heatmap showed the spatial proximity of annotated cell types. Neighborhood enrichment of the whole section. The radius threshold was set to be 50µm. **F.** Representative CNs and neighborhood enrichment result of CN0, CN1 and CN5 respectively.

To confirm that outstanding spatial features can be spotted by CN, we quantified the spatial aggregation of annotated cell types by neighborhood enrichment^15^. We observed the aggregation of T cells, macrophages and non-immune cells in whole slide (Fig 2E). The enrichment analysis of individual CN showed that each CN had a dominant cell type with a spatial aggregation (Fig 2F), which was increasingly obvious than that in whole slide (Fig 2E). Aggregation of neutrophils can be observed in both analyses, though, CN based analysis suggested that the aggregation had a space preference in CN5. The aggregation of Teff in CN0 was ignored in whole slide based analysis, which might be resulted from small portion of Teff and predominantly transcriptomic expression of other cells. Therefore, it was at high risk to neglect the important candidates for in-depth analysis without spoting CN of interest before molecular analysis. In the coming section, we would like to demonstrate the key module of StereoSiTE, SCII.

### Spatial Cell Interaction Intensity (SCII) : Inferring spatially cell-cell communications

The association of cellular distance with cell-cell communication is proposed by stimulated experiment and computational calculation^16^. To validate the impact of distance on communicative activities, and establish recommended parameters, we performed an evaluation procedure to compare results from SCII at different distance thresholds.

To reduce the variance among open-sourced L-R databases, we unified L-R database in SCII by choosing L-R dataset in CellChatDB^5^, which assigned each L-R with an interaction distance associated classification as secreted signaling, ECM receptor and cell-cell contact. Fig 3A demonstrated the count and overlap of inferred cell-cell communications detected by SCII at radius of 50µm, 100µm and 200µm and by CellPhoneDB in CN0 region, which contained 131013 cells. The proportion of interaction types (Fig.3B) at different distance thresholds indicated the cell interactions associated with distance. To be noted, the inferred L-R results had limited overlap between SCII and CellPhoneDB despite of distance thresholds. We calculated the median cell distance of communications which were detected by Squidpy alone and by both Squidpy and SCII (Fig 3C). The distances between neighbored cell pair involved in communications exclusively predicted by CellPhoneDB were significantly longer than others, indicating limited reachable interaction. In other words, CellPhoneDB detected many false positive interactions, which could be avoided in SCII by including distance threshold in analysis.

**Figure 3.**
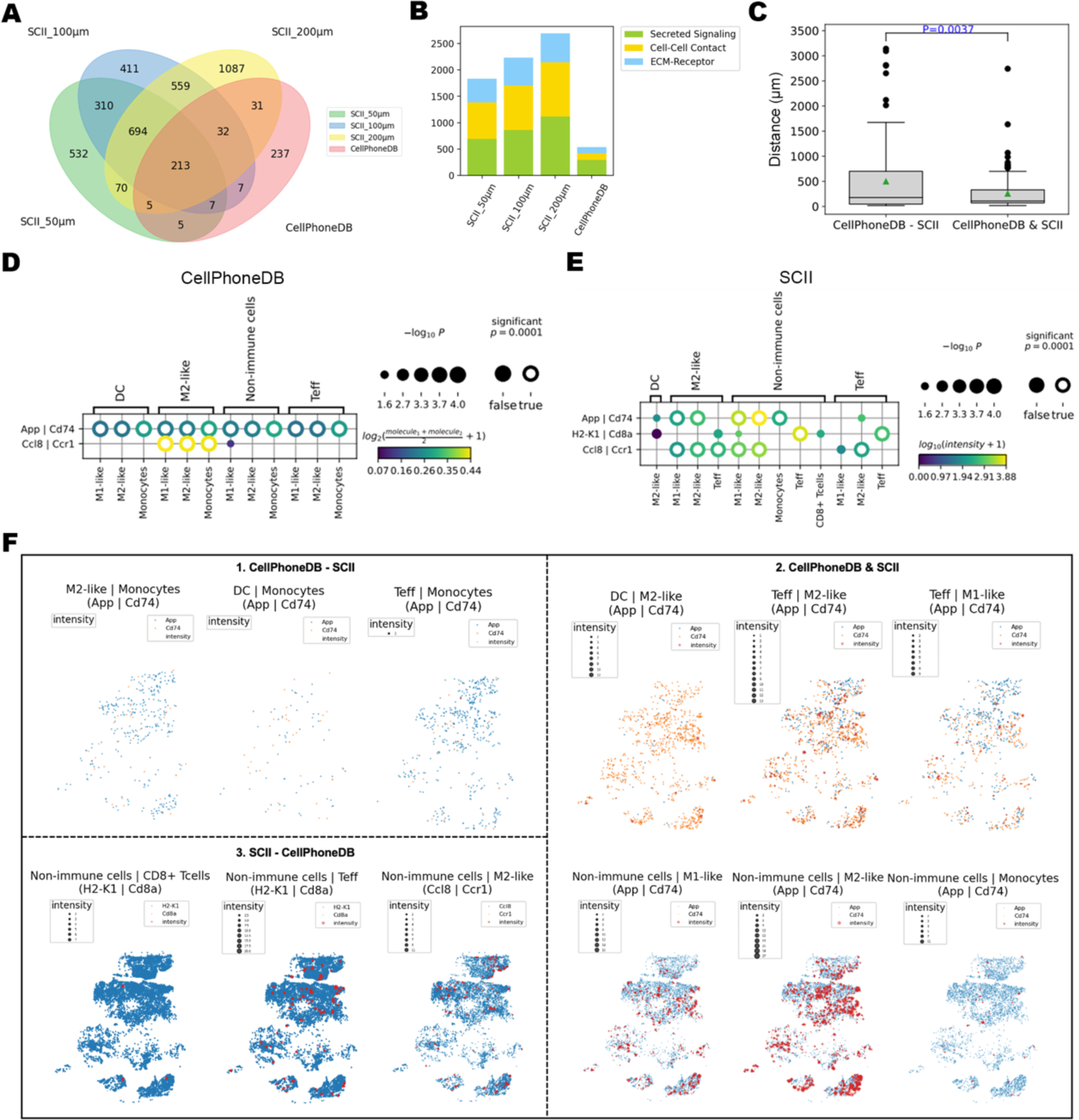
Superior performance of Spatial Cell Interaction Intensity (SCII) compared to analysis without spatial inference. **A.** Venn diagram showed the intersection and difference in predictive interactions from SCII with radius threshold of 50µm, 100µm and 200µm, and from CellPhoneDB. **B.** Stacked bar plot showed the number of predictive interactions of different types detected at radius threshold of 50µm, 100µm and 200µm. Green indicated secreted signaling, yellow indicated cell-cell contact and blue indicated ECM-receptor. **C**. For each communication, we connected sender cell with its N surrounding receiver cells to construct space nearest neighbor graph by KNN, then calculated the median distance of all connected cell pairs. We calculated the median cell distance quantile of communications detected by CellPhoneDB alone and by both CellPhoneDB and SCII. Shown is an approx one-sided P value from Wilcoxon rank-sum test. The green triangle indicates mean value. **D.** The representative results predicted by CellPhoneDB. The color of the bubble indicated the average expression of the L-R pairs, and the size indicated confidence. The confidence p-value less than 0.0001 was indicated by circle. **E.** The representative results predicted by SCII. The color of the bubble indicated the intensity strength, and the size indicated the same as D. **F.** Communications between sender cells and receiver cells mediated by corresponding ligand-receptor interaction were displayed in situ. 1. CellPhoneDB - SCII: interactions detected by CellPhoneDB alone. The intensity showed a low possibility of active interaction between predictive L-R pairs in corresponding cells. 2. CellPhoneDB & SCII: interactions detected by both CellPhoneDB and SCII. The intensity showed a high possibility of active interaction between predictive L-R pairs in corresponding cells. 3. SCII – CellPhoneDB: interactions detected by SCII alone. The intensity showed a strong possibility of active interaction between predictive L-R pairs in corresponding cells. Blue spots indicated sender cells with ligand expression, orange spots indicated receiver cells with receptor expression, red spots indicated local interaction intensity.

Fig 3D and 3E showed representative L-R pairs identified by CellPhoneDB and by SCII, respectively. We mapped the intensity of inferred cell-cell communications in situ in Fig 3F. Unlike exclusively predicted by CellPhoneDB, the L-R pairs identified exclusively in SCII and both in CellPhoneDB and SCII, had stronger spatial interaction (in Fig 3F). The interaction induced by App and Cd74^17^ need direct contact of sender and receiver cells, demonstrating that the introducing distance threshold could avoid false positive caused by unreachable cells. Fig 3F showed the communications exclusively detected by SCII between non-immune cells with CD8+ Tcells, Teff and M2-like could be observed with high intensity of interaction, which supported the superior accuracy of SCII over methods without considering spatial information.

### Profiling tumor microenvironment using spatial transcriptomics

To apply the designed framework to address iTME associated research questions, we introduced xenograft models (Fig 4A) with immune agonist (STING agonist) treatment^18^. Here, we collected spatially resolved transcriptomic data from xenograft tumor tissues by Stereo-seq, a spatial sequencing technology with the subcellular resolution of 500nm^19^. Data matrix at resolution of single cell was obtained after cell segmentation processing based on nuclear staining^13^. By cell2location induced deconvolution of spatial transcriptomic matrix with reference previously reported^14^ we identified and validated 12 distinct cell types (Fig 4B), including 6 of lymphoid-lineage, 5 of myeloid-lineage and 1 of non-immune cluster. With proportion of cell types across samples (Fig 4C), we noticed the different compositions of immune cells across samples. Further comparison in quantitative analysis indicated less cell numbers in treatment group when compared to control (Fig 4D). It was proposed that necrosis caused by treatment might attribute to reducing cell numbers in treatment group (Fig 4E). Notably, control groups had higher frequency of M2-like macrophages, meanwhile, treatment groups possessed higher frequencies of neutrophils (Fig 4C). Interestingly, we additionally observed a location preference (Fig 4F) of neutrophils in treatment group, where they tended to cluster around necrosis niches, vice versa, other cells like M2-like macrophages in control group were randomly distributed. However, methods interrogating the correlation between specific bioactivities and the corresponding spatial preference were rarely exploited. To validate our hypothesis of spatial preference between different cell types, we applied analysis of CN in following section.

**Figure 4.**
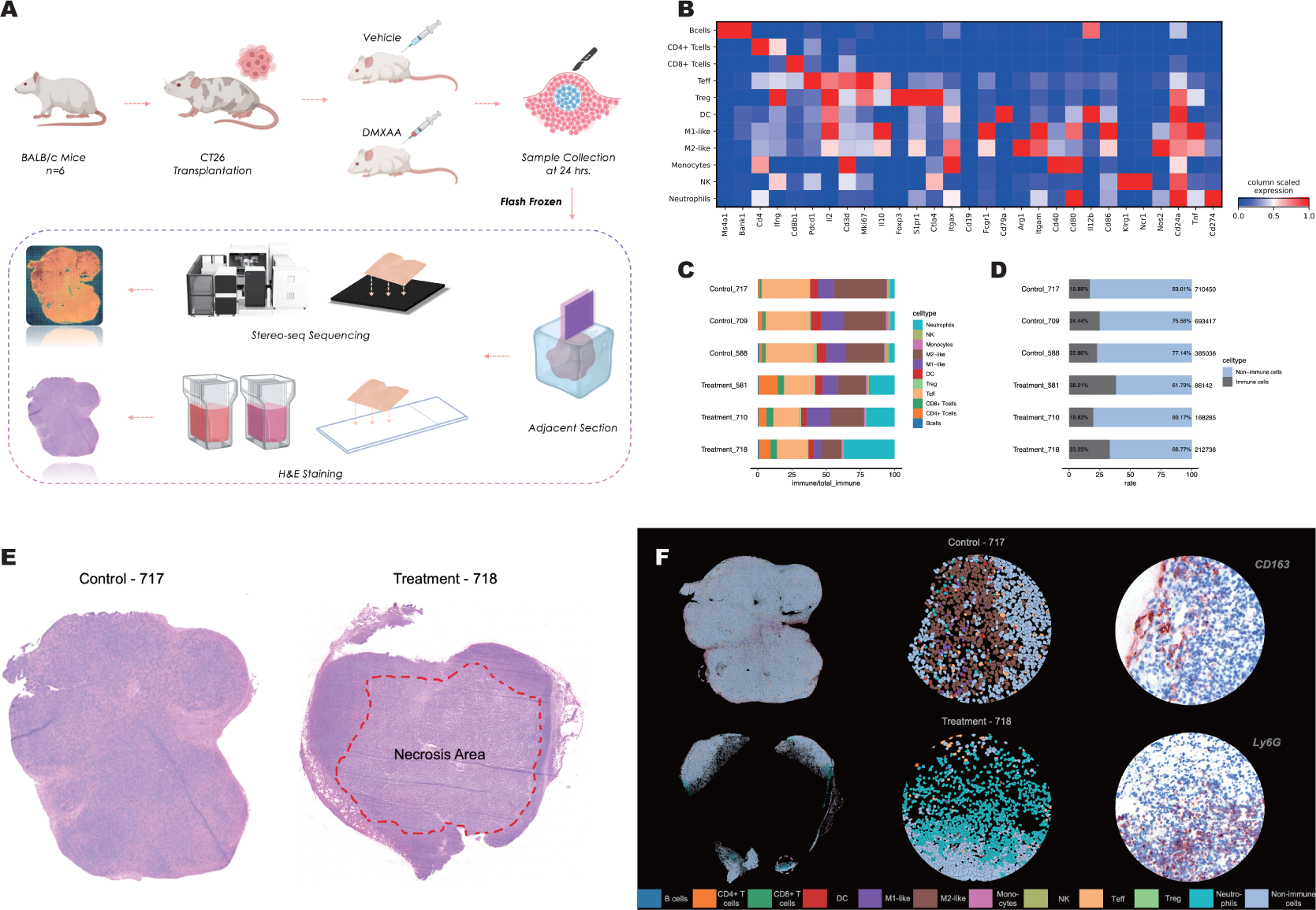
Spatial Transcriptomic mapping of xenograft model in situ. **A.** Flow chart of xenograft model construction. BALB/c mice were subcutaneously injected with colon cancer originated CT26 cells and treated with vehicle or DMXAA for duration of 14 days after tumor transplantation and samples were harvested 24 hours after treatment. **B.** Heatmap of transcriptional markers expression of 11 annotated immune cell types at single-cell resolution. **C.** Analysis of immune cell proportion at single cell resolution in each sample. **D.** Histogram of non-immune cells vs immune cells (blue bar represents non-immune cell proportion and grey bar presents integrated immune cell proportion; numbers displayed aligned to statistical bars indicate precise cell counts regarding to each sample). **E.** Representative H&E staining of sample 717 from control group (left) and sample 718 from treatment group (right). Red-thread restricted region presents necrosis site. **F.** In situ visualization of annotated cell types using Stereo-seq data (Left); Enlarged image of marked-circle-site exhibiting cellular compositions (Middle); Representative IHC staining (CD163 indicating macrophage in sample 717 and Ly6G indicating neutrophils in sample 718) of same marked-circle-site in middle column (Right).

### Organizing TME-associated cellular neighborhood

The immune tumor microenvironment (iTME) heterogeneity prevails both intra- and inter-tumor, which is intrinsically attributed by varied cell organizations in each spatially compartmented unit. The most possible extension to interrogating and elucidating iTME of xenografts, in short of distinct histological characteristics, is to visualize tissues with CNs. With the aim of organizing iTME units that were consistently conserved across samples, we integrated the matrix co-currently covering cellular neighborhoods (CNs) and cell types (CTs) and introduced Tensor to decompose the module matrix^12^. We first clustered windows at ranging size labelling all samples and identified distinct and exclusive CNs under different benchmarks (Fig 5A and Supplementary 1A & 1B). By performing tensor decomposition in different groups (Fig 5B and Supplementary 1C), we observed a given CN correlating with specific CT in each individual module (Fig 5B) and distinct euclidean distance reflecting inter-module heterogeneity (Fig 5C and Supplementary 1D) in the context of bin size 100 micron, we therefore decided to interrogate with this index. As expected, constitution of CTs in each CNs significantly varied across the cohort, which in return suggested that different immune cells preferentially co-localize and associate with certain cell types in compartmented iTME units. We therefore entitled each CN based on their dominant cell proportions (Fig 5D). We next calculated the frequencies of CNs in different groups (Fig 5E) and observed a distinct correspondence of CN3 (NK cell lead), CN4 (Mixed) and CN5 (neutrophils lead) in treatment group (Fig 5F), which was aligned with the treatment background and the tensor indication. To spatially exhibit and evaluate CNs (Fig 5G), we sought to reinforce and assess CN5 in situ since neutrophils were markedly recruited by chemokine motivation but not persistent in targeted tissues^20–22^. We therefore orthotopically projected CNs to adjacent H&E staining to probe the putative distribution pattern of this neutrophils dominant iTME unit (Fig 5H and Supplementary Fig 1E). An obvious trend of CN5 co-localizing around necrosis edges comparing to that of other CNs was observed, which potentially highlighted regional bioactivities exserted by tumor cells after immunoagonist treatment and evidenced alignment between tensor indicated transcriptomic traits and histological features, we therefore decided to further interrogate the bioactivities burst in CN5 in the following work.

**Figure 5.**
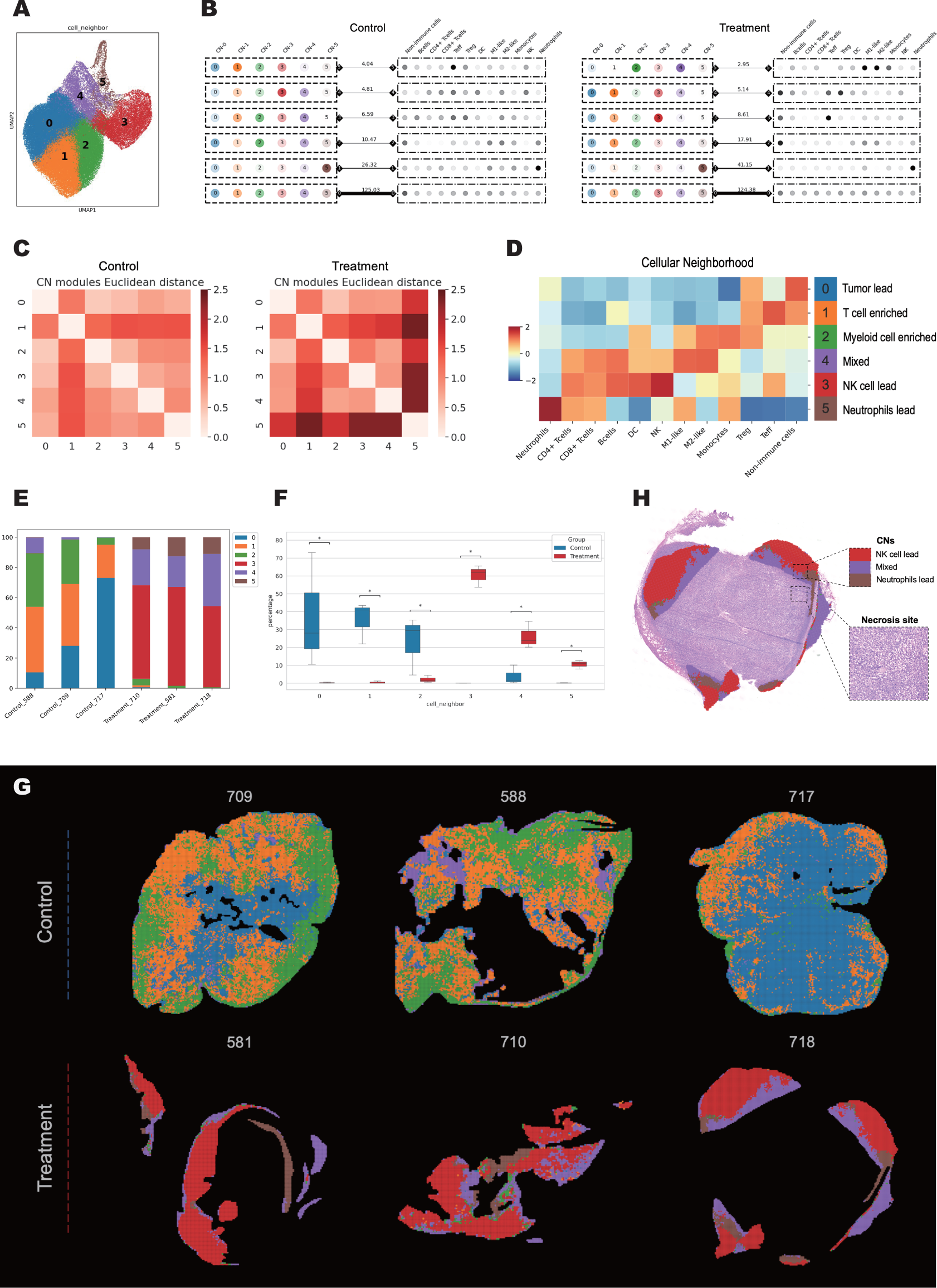
Construction of cellular neighborhood. **A.** UMAP exhibiting the deconvolution of the identified CN clusters at bin size of 100. **B.** Tucker tensor decomposition of control (left) and treatment (right) samples to stratify CN modules and CT modules. The crosstalk extent of associated CN and CT was represented by weight of the line with indicated numbers. **C.** Heatmap of Euclidean distance between CN modules constructed in control (left) and treatment (right) group, respectively. **D.** Heatmap indicating varied cell composition in different CNs (left) and confetti labeling the corresponding title for each CN (right). **E.** CN frequencies in different samples from control and treatment group. **F.** Box plot indicating the statistical variation of CN frequency across groups. **G.** In situ distribution of CNs in different groups. **H.** Projection of CN5 on adjacent H&E staining of sample 718, with enlarged image of marked-circle-site displaying highly resolved H&E staining of necrosis area.

### Deconvolving iTME regarding to individual CNs

To comprehensively depict the distinct landscape STING agonist induced, differentially expressed genes (DEG) analysis was first performed. Each CN was compared to the rest counterparts in treatment group and a potent recruitment of neutrophils and activation of STING signaling was exclusively observed in CN5^22, 23^ (Fig 6A & 6B and Supplementary Fig 2A). In addition, other CNs in treatment group displayed different phenotypes (Supplementary Fig2B), highlighting the drug-response imprinted function in CN5. Since neutrophils were leading in this milieu, we next decided to decipher the signature activities of CN5 at cellular level, SCII was applied to infer the cell-cell communications at single cell resolution. The result of CN5 indicated a potent interaction between non-immune cells and neutrophils cells with representative pair like Cxcl1/Cxcr2 and synchronously revealed prominent neutrophils-neutrophils communication with overload L-R pairs expression like Il1b/Il1r1 and Cxcl2/Cxcr2, (Fig 6C and Supplementary table 1). Meanwhile, the in situ intensity visualization of Cxcl1/Cxcr2 projecting on adjacent H&E image revealed a frequent crosstalk between non-immune cells and neutrophils around necrosis niches (Fig 6D and Supplementary Fig 2B). To deeply investigate the L-R pairs associated to treatment response, we explored the functional signaling pathways in CN5. By performing in silico identification of active transcription factors and potential interactions (Fig 6E), we identified an Nf-kappa b-centric and an Irf1-centric network coordinating the upregulation of signature downstream targets respectively like Il1b, and Ifnβ (Fig 6F). Interestingly, Irf1 was herein determined responsible for Ifnβ regulation while Irf3 was hardly activated at this stage of treatment (Supplementary Fig 2D). Protein-protein interaction (PPI) analysis was next performed to construct the signaling network and to investigate the hub genes responsible for the signature activities in CN5. Excitingly, Il1b was the top scored gene frequently interacting with other proteins, including Ccl4, Cxcl2, Cxcl1 and Il6 (Fig 6G), which was the acquiescent downstream target of NF-kappa B in the context STING signaling activation^23^. The analysis of TF regulon and PPI network coordinately suggested that neutrophils might play key roles of regulating immunoactivities in response to STING agonist via signature signaling pathways like Nf-kappa-b and Irf1. Altogether, we provided an integrated analysis flow to depict the functional iTME unit at both molecular and cellular level.

**Figure 6.**
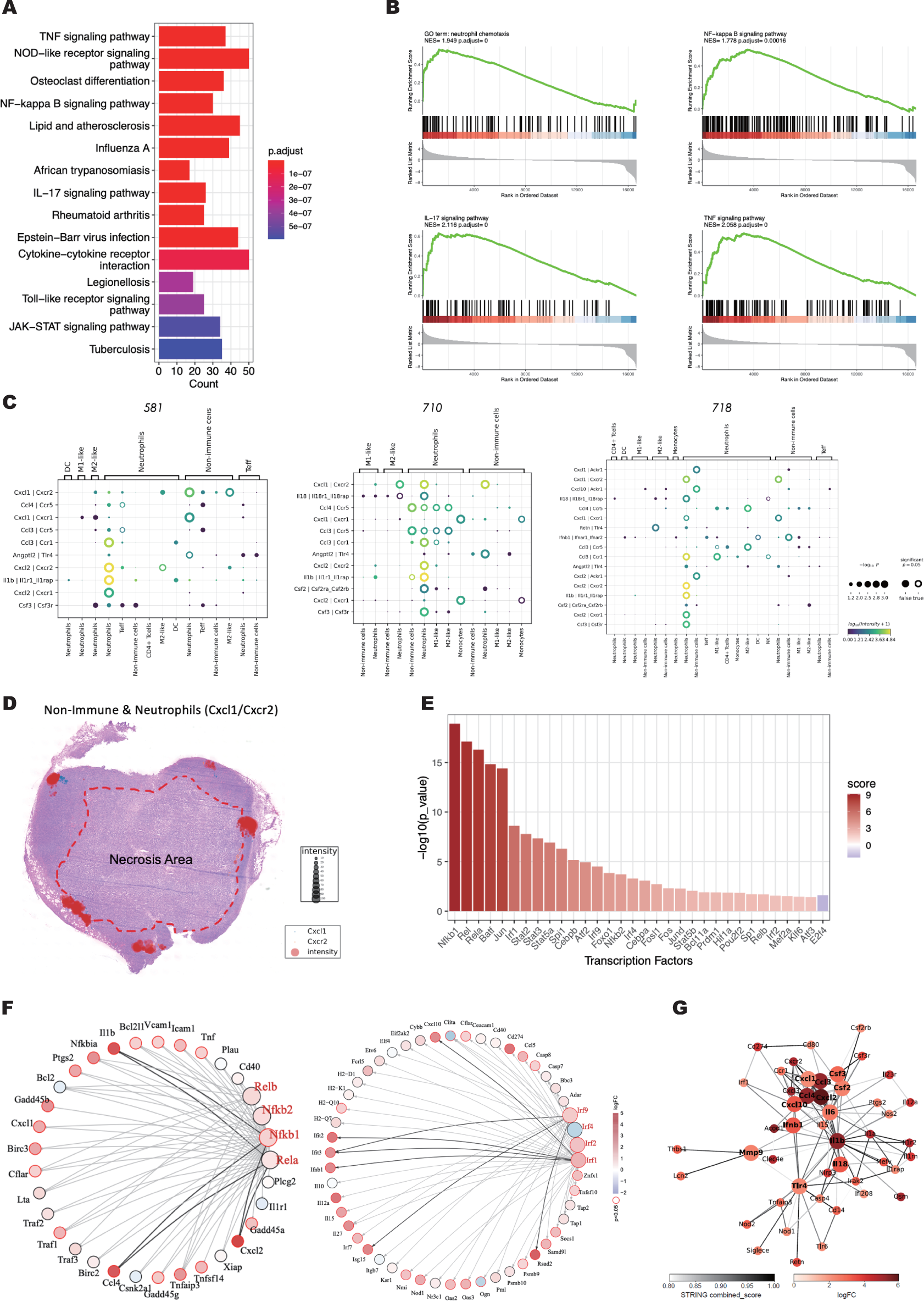
Deconvolution of CN of interest. **A.** KEGG analysis of upregulated signaling pathways in CN5 comparing to that in counterpart CNs. **B.** GSEA analysis of indicated signaling in CN5 comparing to that in counterpart CNs. **C.**SCII calculation of representative ligand-receptor interaction in CN5 of each sample from treatment group. **D.** In situ visualization of Cxcl1/Cxcr2 between non-immune cells and neutrophils with projection to adjacent H&E image to exhibit the crosstalk coordinates. **E.** TF regulon analysis of CN5, score represents the potency of transcriptional factor activity. **G.** Graph indicating predicted interactions between genes differentially (red, transcription factors; black, other genes) expressed in CN 5, (left) exhibited Nfkb1-centric network and (right) exhibited Irf-centric network. **H.** PPI network analysis in CN5 of treatment group, chroma of nodes represents logFC of indicated gene expression and edges represents STRING database score (confidence level). Hub genes were presented with bold font and larger size.

## Discussion

With spatial transcriptomics data at single-cell resolution available^19^, The state-of-art sequencing method enables researchers to obtain the genuine landscape of TME without unbalanced loss of cells caused by dissociation. With multi-dimensional and high throughput of gene expression, it requires powerful algorithmic framework to efficiently associate specific cell types to communicative activities, biological processes and clinical manifestation to address scientific questions^24^. In this paper, we developed a framework called StereoSiTE to identify cellular neighborhood unit harboring disease associated cell types, and to functionally decipher underlying biological activities of signature CN by inferring the active cell-cell communications with supplemented analysis of regional transcriptomic network. StereoSiTE is applicable to spot-based spatial datasets and also compatible with high resolution resolved data like single cell resolution.

Spatially organized iTME and its disorganized manner can be manifested as pathological disease. In StereoSiTE, CN analysis helped to target the critical iTME unit with involved cell types while SCII demonstrated the potential communication activities happened in these functional units with indication of associated L-R pairs. We recommend CN based analysis to identify the regions of interest with indication of neighbored cellular composition which was more likely to have active interaction due to the proximity. CN based analysis also could accurately find the cell types with spatial signature despite low cell numbers and low transcriptomic activities. Moreover, CN based analysis offered the potent candidates for SCII to find out functional cell-cell communications, and promised the exploring of molecular mechanism with PPI and TF network, etc. Referring to scientific questions, StereoSiTE can provide clinicopathogenesis or biological event associated CNs. In previous study, spatial proteomics based CNs provided insights in CRC patients prognosis prediction with specific CNs and cell types^9^, however, the limited throughput dilemma that proteomics-based technology inherent hindered the investigation of regional molecular mechanisms in each iTME units, we therefore designed this framework.

The foundation is the self-developed algorithm, SCII, which infers the active Ligand-Receptor pairs with consideration of spatial distance between interacting cells. Based on this consensus, SCII constructs the graph by connecting cell pairs within a defined radius threshold. The radius threshold could be set referring to communication types. Radius threshold is recommended to be set as 100µm ∼ 200µm for interactions of secreted signaling, while L-R pairs of cell-cell contact should consider close cellular proximity. Therefore, SCII incorporates the reference from CellChatDB which provides catalog associated to communicative distance. A benchmarked analysis is performed with CellPhoneDB and SCII with the same Stereo-seq data matrix. Figure 3 highlights that the analysis without distance threshold may lead to false positive results. On the contrary, enriched L-R pairs in a CN of interest within distance threshold possess high potential for association to functions.

To prove the overwhelmed performance of SCII, we applied the framework in a research scenario to explore the functional cell-cell communications in response to immuno-agonist treatment. The downstream targets and activated pathways coordinately suggested that neutrophils were markedly recruited to CN5 and STING signaling were distinctly activated in this unit. Further SCII analysis revealed a frequent crosstalk between neutrophils and neutrophils/non-immune cells and subsequent in situ mapping implied a location preference of certain L-R pairs, like Cxcl1/Cxcr2. The TF analysis and PPI network analysis were next performed and indicated the most contributing L-R pairs, which were the downstream targets of STING activated signaling – NF-Kappa B^23^. Altogether, we depicted a neutrophil leading iTME units by defining the regional phenotype from both molecular and cellular view, which might shed light on neutrophils exserted mechanism in iTME.

Here, we presented a practical framework (StereoSiTE), which was designed to analyze the spatial transcriptomics data by incorporating open-sourced bioinformatics tools with self-developed algorithm to prove that spatial proximity is a must to guarantee an effective investigation.

## Method and materials

### Mice and cell lines

Female BALB/c mice at 6 weeks of age were obtained from GemPharmatech Co., Ltd. All mice were housed in a specific pathogen–free animal facility at GemPharmatech Co., Ltd. CT26 colon cancer cells were purchased from ATCC. Cells were cultured at 37 oC under 5% CO2 in DMEM supplemented with 10% FBS and 1% penicillin/streptomycin.

### Xenograft tumor models and treatment

We implanted 5×10^5 cells/100μl CT26 cells into the right flanks of BALB/c mice. When the tumors reached a volume of 250-300 mm^3^, we performed intratumorally injections of the STING agonist (0.5 mg/50ul/mouse, DMXAA, Vadimezan) (43). Mice in the control group were intratumorally injected with the same volume of PBS, accordingly. Xenograft tumor samples were collected 24hours after treatment and embedded by OCT on dry ice.

### Stereo-seq library preparation and sequencing

#### Tissue processing

Two consecutive cyro sections of 10 μm were prepared. One section was attached to glass slide and stained by H&E staining following previous protocol. The second section was adhered to the Stereo-seq chip surface and incubated at 37°C for 3-5 minutes. Then, the sections were fixed in methanol and incubated for 40 minutes at −20°C. Stereo-seq library preparation and sequencing followed previous published protocol^19^.

### In situ reverse transcription

Prepared section was processed according to the Stereo-seq Transcriptomics Set User Manual (STOmics) and all reagents were from the Stereo-seq Transcriptomics T kit and Stereo-seq Library Preparation kit (STOmics). Briefly, after washed with PR rinse buffer, tissue sections placed on the chip were permeabilized at 37°C for 10 minutes. RNA released from the permeabilized tissue and captured by the DNB was reverse transcribed overnight at 42°C. After reverse transcription, tissue sections were digested with Tissue Removal buffer at 55°C for 30 minutes. The resulting cDNA was then amplified.

### Amplification

The collected cDNAs were amplified with KAPA HiFi Hotstart Ready Mix (Roche, KK2602) with 0.8 μM cDNA-PCR primer. PCR reactions were performed in sequential steps as incubation at 95℃ for 5 minutes, 15 cycles at 98℃ for 20 seconds, 58℃ for 20 seconds, 72℃ for 3 minutes and a final incubation at 72℃ for 5 minutes.

### Library construction and sequencing

The concentrations of the PCR products were quantified by Qubit™ dsDNA Assay Kit (Thermo, Q32854). A total of 20 ng of DNA were then fragmented with in-house Tn5 transposase at 55°C for 10 minutes. The reactions were stopped by the adding of 0.02% SDS and gently mixing at 37°C for 5 minutes. Fragmented products were amplified as follows: 25 μl of fragmentation product, 1 × KAPA HiFi Hotstart Ready Mix and 0.3 μM Stereo-seq-Library-F primer, 0.3 μM Stereo-seq-Library-R primer in a total volume of 100 μl with the addition of nuclease-free H2O. The reaction was then run as: 1 cycle of 95°C 5 minutes, 13 cycles of 98°C 20 seconds, 58°C 20 seconds and 72°C 30 seconds, and 1 cycle of 72°C 5 minutes. PCR products were purified using the AMPure XP Beads (0.6× and 0.15×), used for DNB generation and finally sequenced on MGI SEQ-2000 sequencer.

### Data analysis

#### Raw sequencing data analysis

Fastq files were generated by MGI SEQ-2000 sequencer. ssDNA image stitching, tissue cut, gene expression register, and genome mapping, gene counts were performed by online analysis Platform: Stereo Analysis Platform (SAP, https://uat.stomics.tech/sap/researchProject/index.html). Cell segmentation was performed according to published method^13^. Expression profile matrix was divided into non-overlapping bins covering an area of 100 × 100 DNBs (bin100) for further cellular neighborhood construction and functional enrichment analysis. Gene expression matrix for each cell were obtained according to spatial coordinates of cell segmentation result. Then data structure were constructed by Scanpy^25^ in python 3.9 for further analysis.

### Cell type annotation

We used a single-cell transcriptomics dataset of mouse colon cancer cell line CT26^26^ as a reference to deconvolute a mixture of 11 immune cell types and non-immune cells in our Stereo-seq data by Cell2location^10^ with hyperparameter N_cells_per_location= 1, detection_alpha=20. The cell type with maximum abundance was assign to each cell, then cell types frequency was calculated and visualized by R package ggplot2.

### Cellular neighborhood construction

The tissues were binned with side-by-side windows, each with an area of 100 × 100 DNBs (bin100), represented a square with side length of 100 DNB (the unit is capturing site). As the distance between two neighbor sites was 500 nm, bin100 corresponded square with side length of 50µm. Then the cellular composition of each bin100 was deconvoluted by mapping gene expression profile of each cell type from single-cell transcriptomic dataset^14^ to spatial data with cell2location. According to the deconvoluted cell composition matrix, the windows of all samples were subsequently clustered to 7 cellular neighborhoods (CNs) by using KNN graph and Leiden with n_neighbors=19, resolution=0.32. Therefore, the windows with similar cell composition were gathered to form a microenvironment. For each CN, the cell type abundance of all windows in this CN region were summed to calculate percentage and visualized by Python module Seaborn.

### Tensor decomposition

For each group, we constructed a tensor with 3×7×12 dimensions (3 samples, 7 CNs and 12 cell types). We performed Non-negative Tucker decomposition by Python package Tensorly^14^. By calculating the decomposition losses of different combinations of the number of CN modules and CT modules, we selected the suitable rank in the elbow point to perform non-negative tensor decomposition (Figure 5A). The visualization of decomposition result refers to Schürch’s article^9^.

### Functional enrichment analysis and transcription factors activity inference

We performed differential expression analysis on different CNs using the edgeR package^27^ in pseudobulk manner^28^. Differential expressed genes were retained when abs(logfoldchanges)>1 and pvalue<0.05. The KEGG enrichment analysis, gene ontology enrichment analysis and GSEA were employed to dissect the biological function of CN5 using functions of R package ClusterProfiler^29^. The top 15 significantly enriched pathways of KEGG enrichment analysis and gene ontology enrichment analysis were displayed as bar plot, respectively. The interesting pathways of GSEA were shown through the gseaplot2 function. The R package decoupleR^30^ and DoRothEA^31^ was used to perform transcription factors (TF) activity inference with Univariate Linear Model, the most significantly TF regulatory network was exhibited by igraph package.

### Spatial cell interaction intensity

Firstly, we constructed the space nearest neighbor graph based on spatial coordinate of all cells, and cell pairs with distance less than the radius threshold were connected by edges. Secondly, the edges were assigned with weight by summing up the ligand and receptor gene expression of the sender and receiver cells (ends nodes of the edge), then the edges with weight 0 (no co-expression) were removed and the edges with weight more than 0 (co-expression) were reserved. Thirdly, we computed the locally spatial cell interaction intensity of every sender cell with its linked receiver cells by summing the edge weights between them, defined as

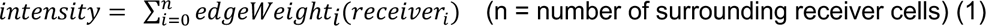

Besides, the interaction intensities from sender cell to receiver cell mediated by specific L-R pairs of the entire slide equaled to the sum of weight of all edges. The formula below presented the computation rule.

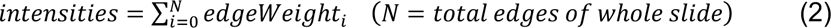

As to the complex ligand or receptor which was composed of several subunits, we selected the minimal expression or calculated the mean expression of all submits to compute the SCII.

### Protein-protein interaction analysis

We queried 628 significant up regulated genes (logFC>2 & FDR<0.05) in CN5 area of treatment samples in STRING v11.5 database^32^ (score cutoff = 0.4) and 505 proteins be found, after filtered 95 nodes with 0 degree, we get a protein-protein interaction (ppi) network with 402 proteins and 2042 edges. Then we performed markov clustering by MCL cluster algorithm^33^ with parameter (I=3.0) to get functional ppi network. The largest cluster with 82 proteins and 821 edges was further used to found hub genes by ranking degree and Maximal Clique Centrality (MCC) score in python module NetworkX 3.1^34^. The highest MCC-scored genes within top 10 degree were labeled as hub genes and their interactions with STRING confidence score over 0.8 were considered as key ppi networks.

## STAR

|  |  |
| --- | --- |
| Stereo Analysis Platform (SAP) | https://uat.stomics.tech/sap/researchProject/index.html |
| Python 3.9 | https://www.python.org/ |
| Numpy 1.22.4 | https://numpy.org/ |
| Pandas 1.5.1 | https://pandas.pydata.org/ |
| Sklearn 1.0.1 | https://scikit-learn.org/ |
| Cell2location 0.1 | https://cell2location.readthedocs.io/en/latest/ |
| Scanpy 1.9.1 | https://scanpy.readthedocs.io/en/stable/index.html |
| Tensorly 0.7.0 | http://tensorly.org/stable/index.html |
| Seaborn 0.11.2 | https://seaborn.pydata.org/index.html |
| Squidpy 1.1.2 | https://squidpy.readthedocs.io/en/stable/index.html |
| NetworkX 3.1 | https://networkx.org/ |
| STRING v11.5 | https://string-db.org/ |
| mcl 22-282 | https://github.com/micans/mcl |
| R 4.2.1 | https://www.r-project.org/ |
| ggplot2 3.4.0 | https://ggplot2.tidyverse.org/ |
| ClusterProfiler 4.6.0 | https://bioconductor.org/packages/release/bioc/html/clusterPr<br>ofiler.html |
| edgeR 3.40.2 | https://bioconductor.org/packages/release/bioc/html/edgeR.ht<br>ml |
| Dorothea 1.10.0 | https://bioconductor.org/packages/release/data/experiment/ht |
|  | ml/dorothea.html |
| decoupleR 2.4.0 | https://www.bioconductor.org/packages/release/bioc/html/decoupleR.html |
| igraph 1.4.3 | https://r.igraph.org/ |

## Conflict of Interest

The authors report no conflicts of interest in this work.

## Acknowledgments

The authors would like to acknowledge China National GeneBank. The authors would like to acknowledge Ms. Meisong Yang (BGI Research-Shenzhen) for her assistance in Stereo-seq experiments.

---

**Supplementary Figure 1.**
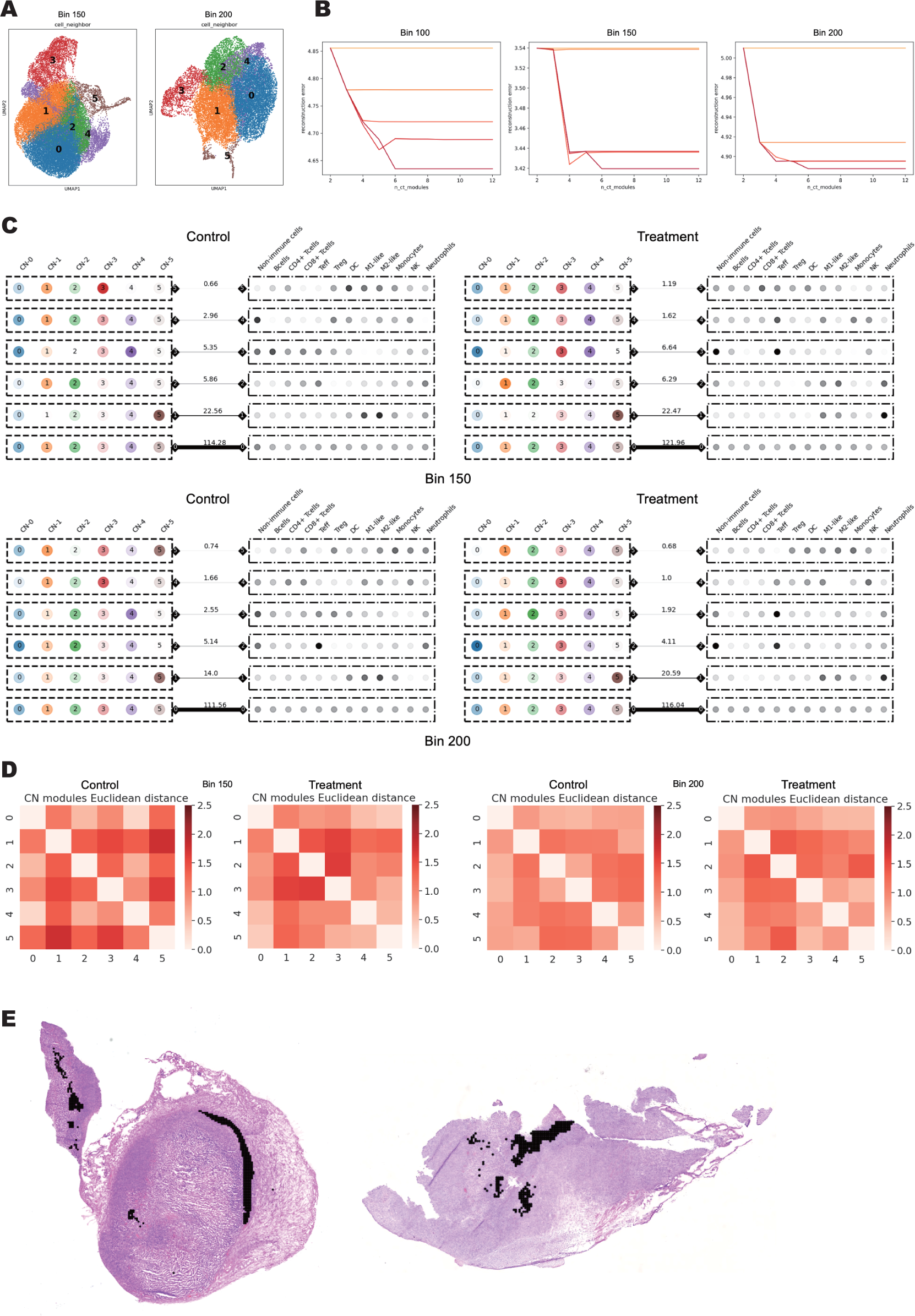
Construction of cellular neighborhood. **A.** UMAP exhibiting the deconvolution of the identified CN clusters at bin size of 150 (left) and 200 (right); **B.** Rank selection of Tucker tensor decomposition at different bin size to stratify CN modules and CT modules. Tensor decomposition loss in different CN modules (different colors) or CT modules numbers (x axis). **C.** Decomposition results for both groups at bin size 150 and 200. The crosstalk extent of associated CN and CT was represented by weight of the line with indicated numbers. **D.** Heatmap of Euclidean distance between CN modules constructed in control (left) and treatment (right) group at indicated bin size, respectively. E. Projection of CN5 on adjacent H&E staining of sample 518 (left) and 710 (right).

**Supplementary Figure 2.**
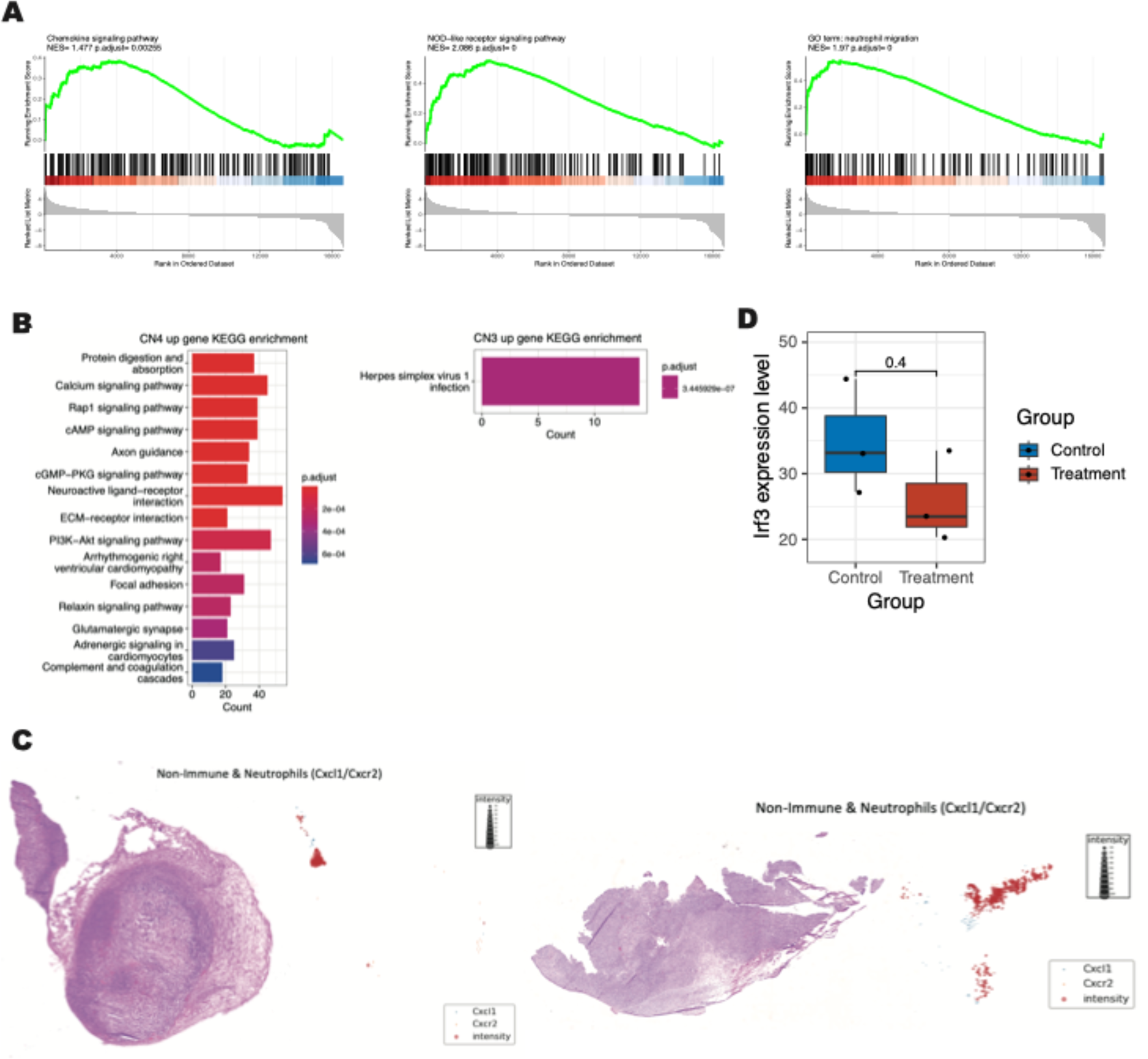
Deconvolution of CN of interest. **A.** GSEA analysis of indicated signaling in CN5 comparing to that in counterpart CNs. **B.** KEGG analysis of upregulated signaling pathways respectively in CN4 (left) and CN3 (right) comparing to that in counterpart CNs. **C.** In situ visualization of Cxcl1/Cxcr2 between non-immune cells and neutrophils in 518 (left) and 710 (right) with adjacent H&E image displayed to exhibit the crosstalk coordinates. D. Box plot of Irf3 expression in control groups and treatment groups (P = 0.4).

## Notes

### Competing Interest Statement

The authors have declared no competing interest.

### Summary of Updates

An update of optimized framework and analytic procedure with better context depiction is made with this revision

